# A SNARE-like mechanism mediates bacterial outer membrane exchange

**DOI:** 10.1101/2025.04.25.650704

**Authors:** Carolina Basurto De Santiago, Elias J. Topo, Pengbo Cao, Daniel Wall, Beiyan Nan

## Abstract

Kin recognition, the ability to distinguish self from nonself at the cellular level is critical to multicellular life. *Myxococcus xanthus* is a multicellular bacterium that cooperates among genetically-related cells and reduces exploitation by nonkin through outer membrane exchange (OME) of common goods and toxins. The polymorphic cell surface receptor called TraA and its partner protein TraB mediate kin recognition by OME, but its molecular mechanism remains unknown. Here we show that TraAB induce OME by overcoming the repulsive interaction between cells. Using quantitative microscopy techniques, we determined that TraA and TraB form complexes at a 1:1 ratio and that as few as one intercellular TraAB-TraAB dimer is sufficient to induce OME. We visualized the OME of single protein particles between cells and revealed that OME depends on the free diffusion of outer membrane (OM) contents. Our findings suggest a mechanism that shows analogies to the eukaryotic soluble N-ethylmaleimide-sensitive factor attachment protein receptors (SNAREs), which mediate plasma membrane fusion. *M. xanthus* OME provides a novel pathway that leads to an underlying conserved mechanism for membrane fusion that is a foundation process for multicellularity.

## Introduction

The ability to distinguish self from nonself at the cellular level is critical to multicellular life. This process ensures that highly related or clonal cells interact cooperatively while excluding unrelated cells, which are often detrimental to multicellularity. For organisms that build multicellularity by aggregating cells from their environment, self/nonself-recognition, or kin recognition, is of paramount importance for coalescing a cooperative kin collective.

Myxobacteria are a group of aggregative multicellular organisms that possess known systems for distinguishing relatedness at the genus and species levels (Kroos *et al*., 2025). Specifically, *Myxococcus xanthus* uses the type VI secretion system (T6SS) and outer membrane exchange (OME) for this purpose (Vassallo & Wall, 2019, Vassallo *et al*., 2020, Gong *et al*., 2018). The T6SS is used by many Gram-negative bacteria for various types of social conflicts and virulence and is relatively well understood (Cherrak *et al*., 2019). In contrast, OME is restricted to myxobacteria and is not as well studied.

OME self-recognition occurs through homotypic binding between adjacent cells, mediated by their polymorphic cell surface receptor called TraA and its operonic partner, TraB (Pathak *et al*., 2013, Cao & Wall, 2017). Following cell-cell recognition, TraAB catalyzes OME, which involves the bidirectional transfer of different outer membrane proteins (OMPs) and lipids between cells (Nudleman *et al*., 2005, Wall, 2014, Pathak *et al*., 2012). This process is thought to occur by transient outer membrane (OM) fusion, where TraAB acts as a fusogen on each interacting cell (Cao *et al*., 2015, Cao & Wall, 2019a). The TraA protein contains a signal peptide (SP), a variable domain (VD) responsible for recognition specificity, a cysteine-rich repeat region that likely functions as a presentation stalk for the VD, and a MYXO-CTERM sorting tag that is likely lipidated to anchor it to the cell surface (Sah *et al*., 2020, Guo *et al*., 2025). TraB contains an SP, an OM β-barrel, and an OmpA cell-wall-binding domain. Forward genetic screens suggest that TraAB are the only proteins required for OME (Dey & Wall, 2014, Vassallo *et al*., 2021).

OME serves two broad functions: mediating cooperation and discriminating between cells (Sah & Wall, 2020). For cooperation, OME can rescue genetic defects in recipient cells by transferring a wild-type protein from donor cells (Hodgkin & Kaiser, 1977, Pathak *et al*., 2012). OME can also rescue lethal OM defects by transferring lipopolysaccharides (LPS) from healthy donors (Vassallo & Wall, 2016). Additionally, when individual cells coalesce into tissue-like assemblages, genetically identical cells can have distinct life histories that result in different adaptations depending on their age and/or microenvironment exposures. Here, OME facilitates the transition to homogeneous and synchronized cell populations with respect to their OM components (Cao & Wall, 2019b), thus facilitating cooperation. Finally, adaptive traits suited to particular environments can be transferred to naïve siblings via OME (Subedi *et al*., 2024), thus preparing or adapting them against future stresses.

OME also functions to discriminate against nonkin. This occurs by the exchange of dozens of polymorphic lipoprotein toxins that belong to six different families (Dey *et al*., 2016, Vassallo *et al*., 2017, Vassallo & Wall, 2019, Weltzer *et al*., 2025). Therefore, when partnering cells are clonal, they are not intoxicated because they express a cognate set of immunity proteins, which themselves are not exchanged. In contrast, when nonclonal cells fortuitously possess compatible TraA receptors, a second discrimination step occurs, wherein cells are reciprocally poisoned because they lack a complete set of immunity proteins. These steps result in an extremely high level of specificity for self-recognition, ensuring that highly related collectives form for multicellular functions, such as fruiting body development.

Membrane fusion plays a central role in many biological processes in eukaryotic cells, but in bacteria there are few examples and little is known. Here, we sought to address key questions about the molecular mechanism of OME. First, we sought to understand the nature of TraAB multimeric complexes that form upon cell-cell contact and OME engagement. Following the coalescing of receptors, these complexes are readily visualized as foci when using functional fluorescent protein fusions to TraA or TraB. However, the stoichiometry of the intercellular TraA junction and the intracellular TraAB complexes remains unknown. Second, since client OME cargos do not contain specificity factors for transfer, but must be diffusible in a fluid OM, we aimed to measure the fluidity of *M. xanthus* OM. Third, do these foci junctions serve as conduits for the transfer of client proteins, and what trajectories do these proteins follow upon OME? Using photobleaching and single molecule tracking techniques, we addressed these questions and proposed an updated model of OME in myxobacteria.

## Results

### Repulsive interaction between cells obstructs OME

Cells lacking functional TraAB or expressing incompatible TraA variants do not exchange their OM contents (Wei *et al*., 2011). This raises the question; why are TraAB required and physical cell-cell contacts alone are insufficient for OM fusion? To answer this, we investigate cell-cell contacts using cryogenic electron microscopy (cryoEM). To avoid the potential interference from extracellular polysaccharides (EPS), we grew *pilA^—^* cells that do not produce EPS (Yang *et al*., 2010, Zhou & Nan, 2017) in liquid culture, concentrated cells by centrifugation, and captured micrographs of *M. xanthus* cells in close contact, where we found cell groups of various sizes. As OME does not occur in liquid, these cells displayed the character of OMs before fusion (**Fig. 1A**).

**Fig. 1.**
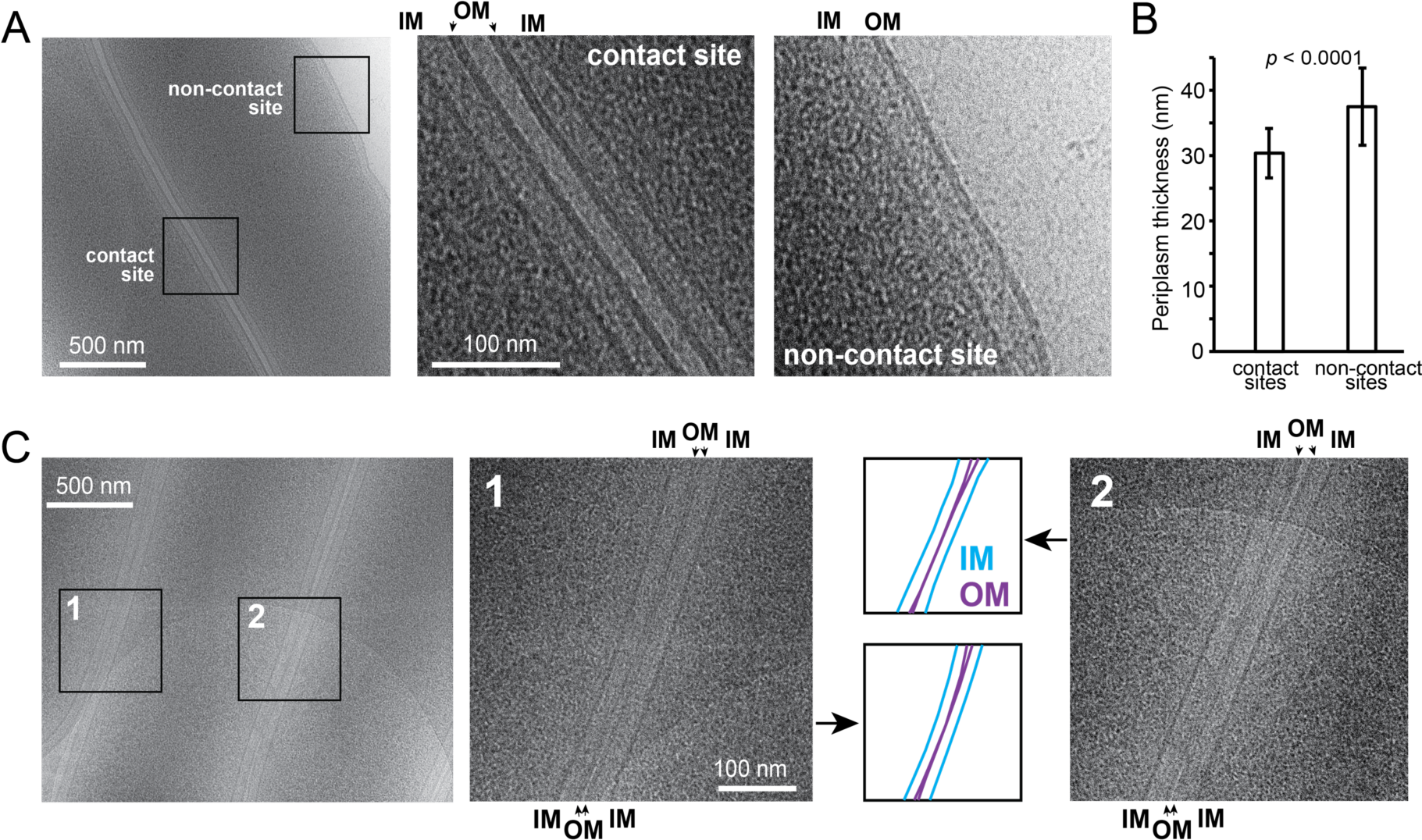
TraAB facilitate OME by overcoming the repulsive interactions between cells. **A)** cryoEM images show that juxtaposed *pilA^—^* cells do not make direct OM-OM contacts. In the same cell, the OM is relaxed at non-contact sites while stressed at cell-cell contact sites. **B)** Cell-cell contact induces stress compression of periplasms at contact sites. Periplasmic thickness was measured as the distance between the centers of the OM and IM densities from seven contacting cell pairs, 30 measurements/pair. The *p* value was calculated using a one-way ANOVA test between two unweighted, independent samples. **C)** Juxtaposed *pilA^—^* cells that over produce TraAB frequently form direct OM-OM contacts that lead to OM fusion.

Analyzing these micrographs, we found two remarkable features. First, regardless of group size, physical interaction between OMs was not observed (**Fig. 1A**). The distances between the centers of the OM densities at side-to-side cell contact sites varied between 20.4 nm and 34.5 nm, with an average distance of 28.8 ± 3.4 nm (measure from three pairs of cells, 30 measurements/pair). Therefore, the interaction between the OMs is under the regime of electrostatic repulsion, likely driven by the strong negative charge on lipopolysaccharide molecules, and beyond the range of van der Waals attraction, 0.3 – 0.5 nm (Batsanov, 2001). Such repulsion is stronger than the one between weakly charged liposomes (3 – 10 nm distance) that prevents their fusion in aqueous media (Jahn *et al*., 2024). Hence juxtaposed cells are always separated by a thin water layer and the repulsion between cells physically obstructs OM fusion. Second, the periplasms appeared compressed at cell-cell contact sites (**Fig. 1A**). Strikingly, in the cells that contact others only on one side, only the contact sides displayed compressed periplasms with the thickness (the distance between the centers of the OM and IM densities) of 30.4 ± 3.8 nm, while the non-contact sides retained relaxed periplasms of 37.5 ± 5.9 nm thickness (measured from the same seven pairs of cells, 30 measurements/pair, for both measurements, **Fig. 1B**). This result shows that cell-cell contacts increase the surface tension of OMs. Therefore, similar to plasma membranes that do not fuse spontaneously (Rand & Parsegian, 1984), repulsion must be overcome to bring OMs closer, which poses a major energy barrier in OM fusion.

### TraAB induce OME by overcoming the repulsive interaction between cells

To visualize how TraAB facilitate OME, we subjected a *pilA^—^* strain that expressed *traAB* ectopically by the strong *pilA* promoter (*traAB^OE^*) (Pathak *et al*., 2012), to cryoEM imaging. In stark contrast to its parent strain that maintained a nearly constant OM spacing (**Fig. 1A**), the OM-OM distance between juxtaposed *traAB^OE^* cells varied considerably. Notably, these cells frequently made close OM contacts that often culminated in an apparent OM fusion-like configuration (**Fig. 1C**). Hence, TraAB-TraAB complexes between cells overcomes repulsive interactions between juxtaposed OMs and thus induces OME.

### Stoichiometry of the intercellular TraA foci

TraAB are thought to function as fusogens to catalyze OM fusion between cells. By using fluorescent fusion reporters to TraA or TraB they were found to coalesce into foci aggregates upon cell-cell contact (Cao & Wall, 2019a). Here we sought to address (i) how many TraA molecules are present in these foci? (ii) Do all the TraA foci contain the same stoichiometry, and (iii) do interacting cells contribute the same number of TraA proteins in each focus? To answer these questions, we used a functional monomeric super folder green fluorescent protein (GFP)-labeled TraA (TraA-GFP) expressed from the heterologous *pilA* promoter and placed it in a Δ*traA* strain (Cao & Wall, 2019a). We mixed this strain with wild-type (unlabeled) cells at a 1:2 ratio and spotted the cell mixture on a 1.5% agar surface. As reported (Cao & Wall, 2019a), TraA-GFP distributed uniformly throughout the OM of isolated cells. However, upon cell-cell contact, we observed two types of TraA foci at contact junctions, bright foci occurred between two labeled cells and dim foci between labeled and unlabeled cells (**Fig. 2A**).

**Fig. 2.**
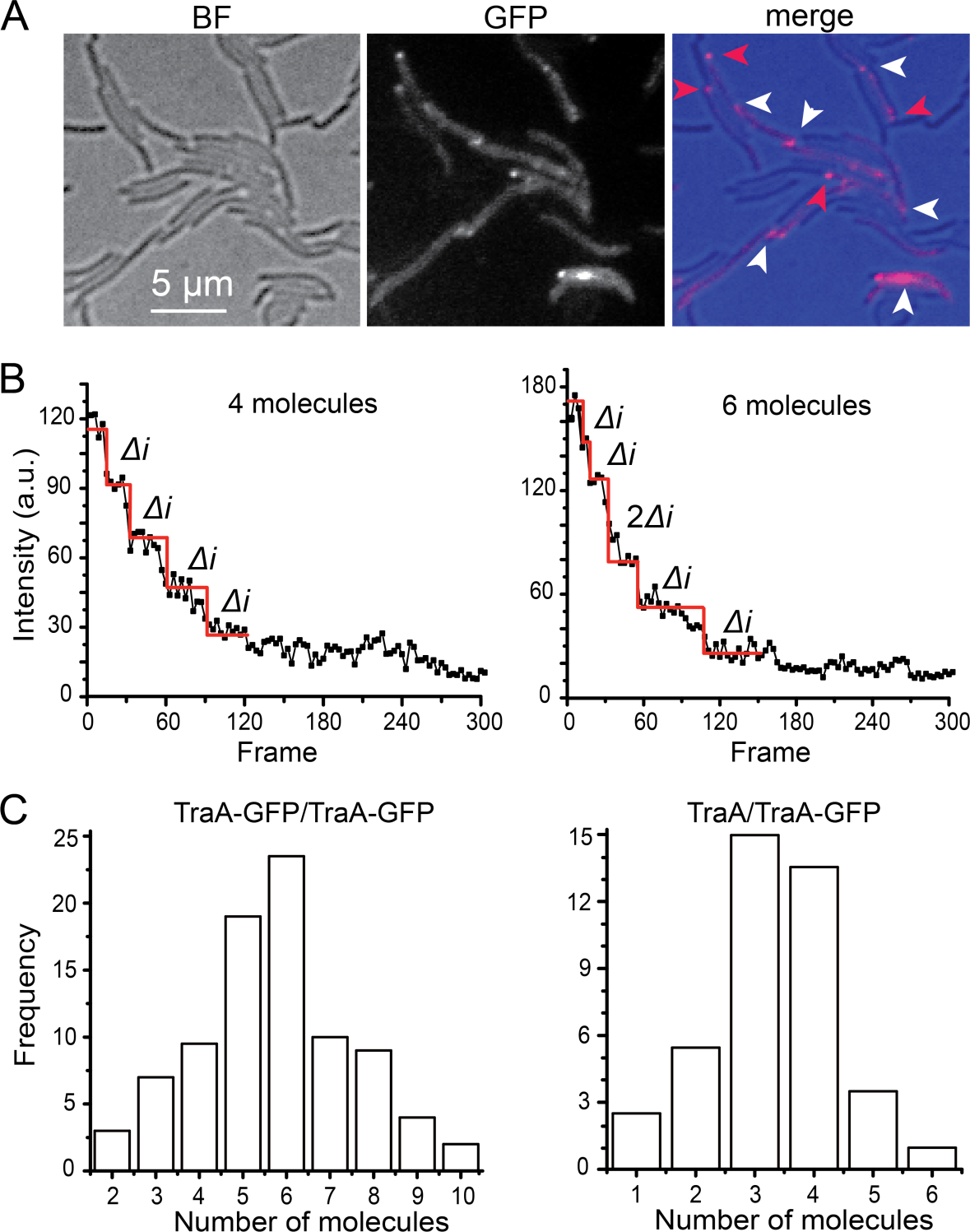
TraA forms heterogeneous foci at cell-cell contact sites. **A)** Representative images of the TraA-GFP foci. The Δ*traA traA-gfp* cells were mixed with wild-type (unlabeled) at a 1:2 ratio. White arrows indicate foci formed between labeled cells and red arrows between labeled and unlabeled cells. **B)** Examples of the photobleached foci that contain 4 and 6 TraA-GFP molecules, respectively. The decrease step of fluorescence corresponds to one GFP molecule being photobleached (*Δi*). **C)** The number of TraA-GFP molecules in each focus between labeled cells (left, n = 87) and between labeled and unlabeled ones (right, n = 41).

The intensities of TraA-GFP foci appeared heterogeneous, regardless of whether only one or both cells were labeled (**Fig. 2A**), suggesting that individual TraA foci do not adopt the same stoichiometry. We then subjected the TraA-GFP foci to photobleaching using a 488-nm laser (0.5 kW/cm^2^) and monitored the decrease of fluorescent signals continuously at 67 Hz (15 ms/frame). The intensity of these foci decreased in discrete steps of the same size, which we define as Δ*i* that reflected the photobleach of one GFP molecule (**Fig. 2B**). Occasionally we saw larger bleach steps where the intensity decreased by 2Δ*i*, indicating that two molecules were bleached simultaneously (**Fig. 2B**). Therefore, the number of Δ*i* bleach steps equal the number of TraA-GFP molecules within the focus (**Fig. 2B, Movie S1**).

Consistent with the variation of their fluorescent intensities, the 87 foci between labeled cells lost fluorescence after 2 to 10 steps of photobleaching, indicating that these foci contained 2 to 10 TraA-GFP molecules (**Fig. 2B, Movie S1**). The numbers of TraA-GFP molecules in each focus displayed a typical Gaussian distribution. Importantly, we never observed any foci that was bleached in a single step of Δ*i*, suggesting that at least one TraA molecule in each juxtaposed cell was required to establish the intercellular junction. This result was consistent with the observation that TraA-GFP expressing cells cannot form TraA foci when it contacts a Δ*traA* cell (Cao & Wall, 2019a). In addition, 46% of these foci contained odd numbers of TraA molecules, indicating that the formation of these foci does not necessarily require the same number of TraA molecules from two juxtaposed cells.

Among the 41 foci found between labeled and unlabeled cells, the number of TraA-GFP molecules also showed a nearly symmetric distribution between 1 and 6, roughly half as many as detected in the foci between two labeled cells (**Fig. 2C**). The detection of foci that contained only one TraA-GFP molecule further confirmed that a single TraA molecule from each contacting cell was sufficient to establish an intercellular TraA junction. Taken together, TraA foci at cell-cell contact sites contain molecules from both juxtaposed cells where each cell may contribute different numbers of TraA molecules.

### TraA and TraB form complexes at a 1:1 ratio

TraA requires its partner OMP TraB to form functional complexes (Pathak *et al*., 2012, Cao & Wall, 2017). Although TraB does not play a specificity role in kin recognition, TraA and TraB colocalize into the foci between adjacent cells in an interdependent manner, suggesting that these proteins form intracellular complexes (Cao & Wall, 2019a). However, the stoichiometry of these complexes is unknown. To address this question, we used a strain that expressed both TraA-mCherry and TraB-GFP and photobleached both mCherry and GFP simultaneously using Hamamatsu W-VIEW GEMINI^TM^ beam-splitter. TraA and TraB colocalized in the foci between juxtaposed cells (**Fig. 3A**). To simplify analysis, we recorded the photobleach of both fluorophores at 5 Hz (200 ms/frame, **Movie S2**). Using this method, we found that the numbers of TraA-mCherry in each focus varied between 2 and 12 in the 59 analyzed foci (**Fig. 3B**), consistent with the photobleach result from TraA-GFP (**Fig. 2C**). Importantly, the TraB-GFP molecules displayed a similar distribution pattern (**Fig. 3B**). We further investigated the correlation between TraA and TraB in each focus and found that each TraA molecule correlated with 0.98 TraB molecules, indicating that TraA and TraB form complexes at a 1:1 ratio (**Fig. 3C**).

**Fig. 3.**
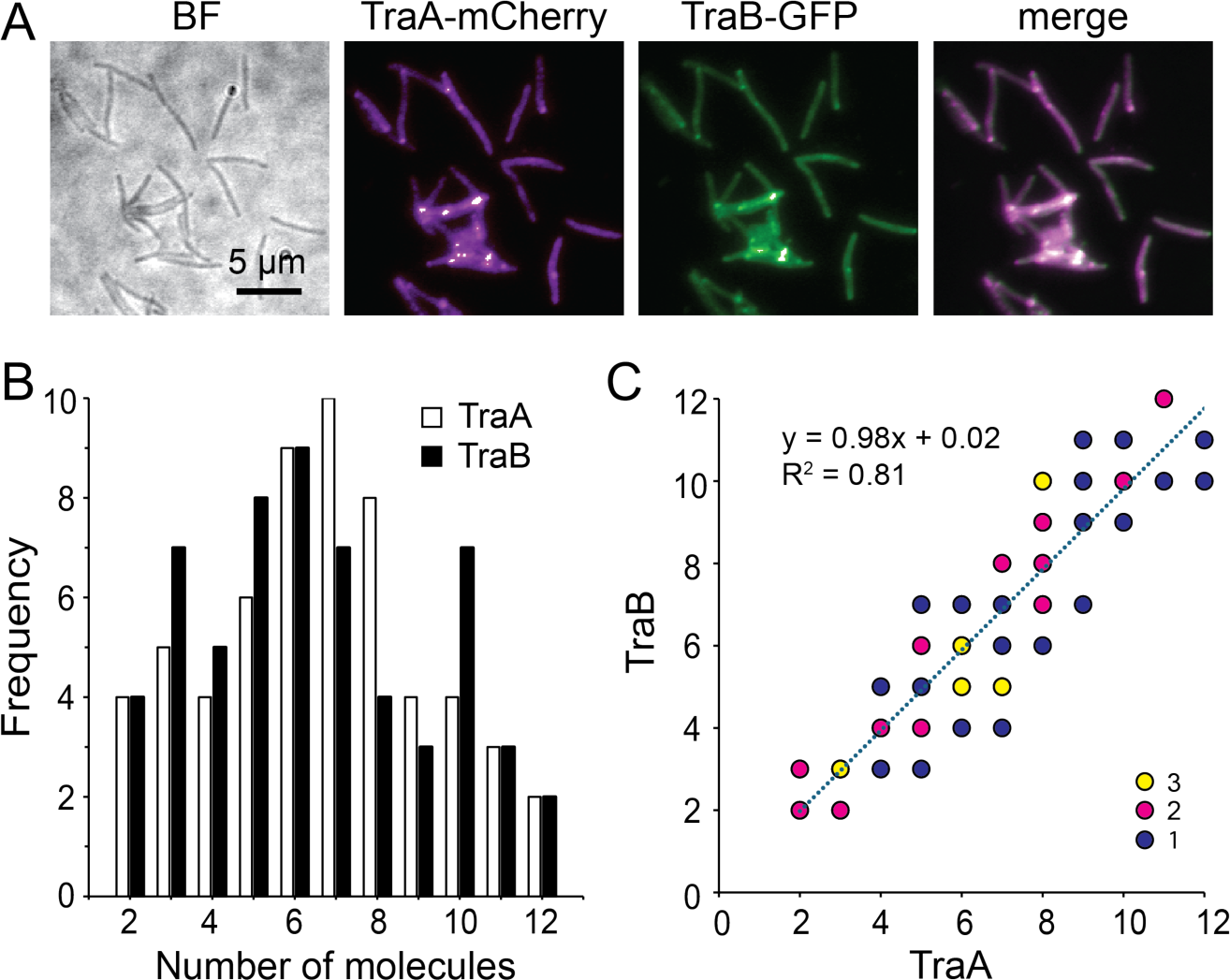
TraA and TraB form intracellular complexes at a 1:1 ratio. **A)** TraA-mCherry and TraB-GFP colocalize in the foci between adjacent cells. **B)** From 59 foci analyzed, the number of TraA and TraB proteins in each focus show similar distribution patterns. **C)** The correlation between the numbers of TraA and TraB in individual foci suggests a 1:1 ratio in the intracellular TraAB complexes. In each focus, the number of TraB is plotted against that of TraA. The numbers of foci that have the same TraA:TraB ratio are indicated by color.

### *M. xanthus* contains a highly fluid OM

The model bacterium *Escherichia coli* has a rigid OM (Rojas *et al*., 2018, Sun *et al*., 2022). As a consequence, the lateral mobility of OMPs is restricted due to the formation of large OMP islands and the selective presence of bulky LPS molecules in the outer leaflet (Rassam *et al*., 2015, Benn *et al*., 2021). In contrast, *M. xanthus* cells are flexible and can exchange a broad range of OM molecules efficiently, including OMPs, LPS, and dyes (Ducret *et al*., 2013, Vassallo *et al*., 2015, Pathak *et al*., 2012). Since heterologous proteins, like a lipidated GFP, are readily exchanged as cargos, OME does not require other *cis* specificity sequences for transfer. Instead, these cargos are likely transferred between cells by diffusion, which requires a fluid OM (Cao & Wall, 2020, Cao & Wall, 2019a). However, the fluidity of *M. xanthus* OM has not been measured.

To measure membrane fluidity, we used the diffusion rate of OMPs as a proxy. Here, we used a lipoprotein where a type II signal sequence was fused to the fluorescent protein mCherry and expressed as a recombinant SS_OM_-mCherry reporter in *M. xanthus.* This simple lipoprotein tag was sufficient for the OM localization of mCherry and enabled it to be transferred by OME (Wei *et al*., 2011). We reengineered this strain by replacing mCherry with the photoactivatable mCherry (PAmCherry). mCherry and PAmCherry are of the same size and share 95.8% protein sequence identity. We used a 405-nm excitation laser (0.2 kW/cm^2^, 0.1 s) to activate the fluorescence of a few labeled SS_OM_-PAmCherry particles randomly (here we refer isolated fluorescence spots as single particles rather than “single molecules” because each spot could contain more than one molecule, albeit only the fluorescence of one molecule was activated). We imaged the activated particles at 67 Hz using a 561-nm laser with total internal reflection fluorescence (TIRF) microscopy (Fu *et al*., 2018, Nan *et al*., 2015, Nan *et al*., 2013, Pogue *et al*., 2018, Ramirez Carbo *et al*., 2024, Zhang *et al*., 2020). Using this setting, only a thin section of each cell close to the coverslip was illuminated, which included the cell surface. To exclude the potential influence of cell-cell contact, we only analyzed the dynamics of SS_OM_-PAmCherry particles in isolated cells. Using single particle tracking photo-activated localization microscopy (sptPALM), we identified 1,518 fluorescent particles that remained in focus for 4 - 12 frames (0.4 - 1.2 s) and plotted their mean squared displacements (MSDs) against time lag (Δ*t*). The linear relationship between MSD and Δ*t* indicated that single SS_OM_-PAmCherry particles moved by typical diffusion (**Fig. 4A, B, Movie S3**). The diffusion coefficient (*D*) of the entire population was 0.19 ± 0.04 µm^2^/s (n = 674, **Fig. 4C**), in the same order of magnitude as the unrestricted diffusion of proteins in the cytoplasm, IM, and periplasm in other bacteria, such as *E. coli* and *Bacillus subtilis* (Parry *et al*., 2014, Lucena *et al*., 2018, Tran *et al*., 2024). Importantly, the diffusion of SS_OM_-PAmCherry was about 10-fold faster than the restricted diffusion of *M. xanthus* proteins in the IM and periplasm (Fu *et al*., 2018, Nan *et al*., 2013, Zhang *et al*., 2023b, Ramirez Carbo *et al*., 2024), suggesting that SS_OM_-PAmCherry diffuses with little restriction in the OM.

**Fig. 4.**
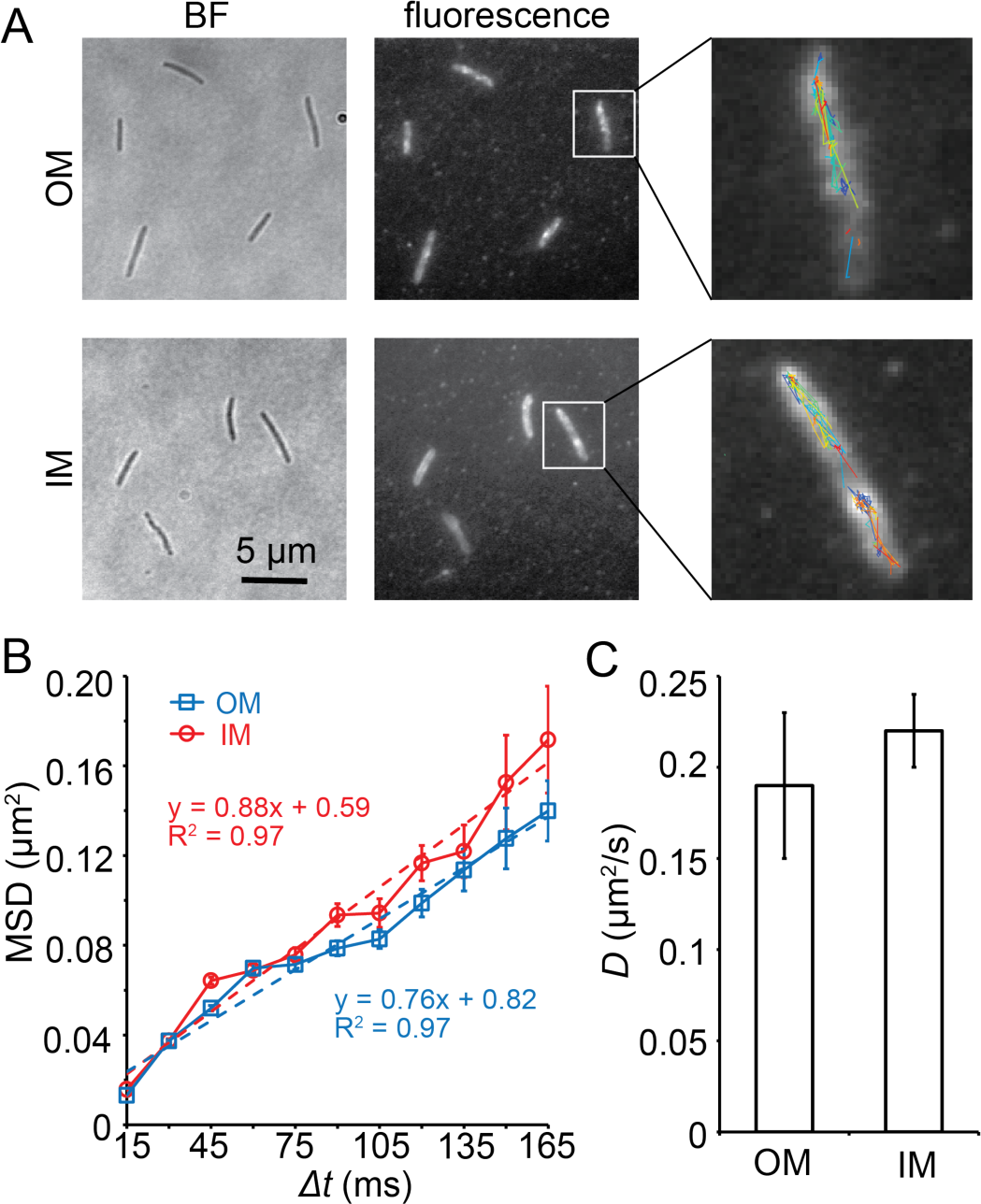
SS_OM_-PAmCherry diffuses rapidly in the *M. xanthus* OM. **A)** Single PAmCherry particles diffuse freely in both the OM and IM. The overall distribution of SS_OM_- and SS_IM_-PAmCherry is displayed using the composite of 100 consecutive frames taken at 15-ms intervals. Single-particle trajectories of PAmCherry were generated from the same frames. Individual trajectories are distinguished by colors. **B)** The linear relationship between MSD and Δ*t* indicates that PAmCherry diffuses freely in both OM (n = 674) and IM (n = 1276). **C)** PAmCherry shows similar diffusion coefficients in OM (n = 674) and IM (n = 1276).

The bacterial IM is known to be fluid, where many proteins diffuse freely (Lucena *et al*., 2018). As a control, we engineered a strain that localizes mCherry to the IM (SS_IM_-mCherry) (Wei *et al*., 2011) by replacing mCherry with PAmCherry. As expected, we found that the recombinant SS_IM_-PAmCherry also diffused freely, with a *D* value of 0.22 ± 0.02 µm^2^/s (n = 1276), close to that of SS_OM_-PAmCherry (**Fig. 4, Move S4**). Taken together, like the IM, the OM of *M. xanthus* is highly fluid and such fluidity provides the mechanistic foundation for OME.

### OME depends on the diffusion of OM components

OME between *M. xanthus* cells has been imaged in real-time (Ducret *et al*., 2013, Cao & Wall, 2019a). However, the mechanism of OME is not fully understood. For instance, does OME depend on the diffusion of the client cargos, or the exchange of large OM fragments? To investigate the dynamics and trajectories of client proteins during OME, we visualized and tracked single SS_OM_-PAmCherry particles. If OME requires diffusion, then SS_OM_-PAmCherry particles should quickly travel between cells. Alternatively, if cells exchange bulk OM fragments, the exchange of SS_OM_-PAmCherry particles should be slow due to the size of such fragments.

To test these possibilities, we used cells that expressed SS_OM_-PAmCherry as donors and Δ*traA* P*_pilA_-traA-gfp* cells as recipients. We mixed these strains at a 1:1 ratio on a 1.5% agar surface. To increase the chance of capturing OME, we first exposed cells under the 405-nm laser for 2 s, where the majority of PAmCherry was photoactivated (Fu *et al*., 2018). We then excited the fluorescence of PAmCherry and GFP using the 561-nm and 488-nm lasers, respectively, and imaged both channels simultaneously using the beam-splitter. As SS_OM_-PAmCherry particles moved rapidly (**Fig. 4**), we had to track their movements at a high frame rate (67 Hz). For this reason, we could only image each field for a few seconds before the reporters’ photobleach (**Fig. 2**). In these short time windows, OME was rarely visualized. Nevertheless, we captured OME of 39 SS_OM_-PAmCherry particles (**Fig. 5A-C, Movie S5-S7**). Their MSD showed a linear relationship with Δ*t*, with a *D* value of 0.21 ± 0.07 µm^2^/s (n = 39), close to the intracellular diffusion rate of SS_OM_-PAmCherry particles. These results indicate that cell-cell contact does not hinder OMP diffusion, and that OME depends on the diffusion of OM content, rather than the transfer of large OM fragments (**Fig. 5D**).

**Fig. 5.**
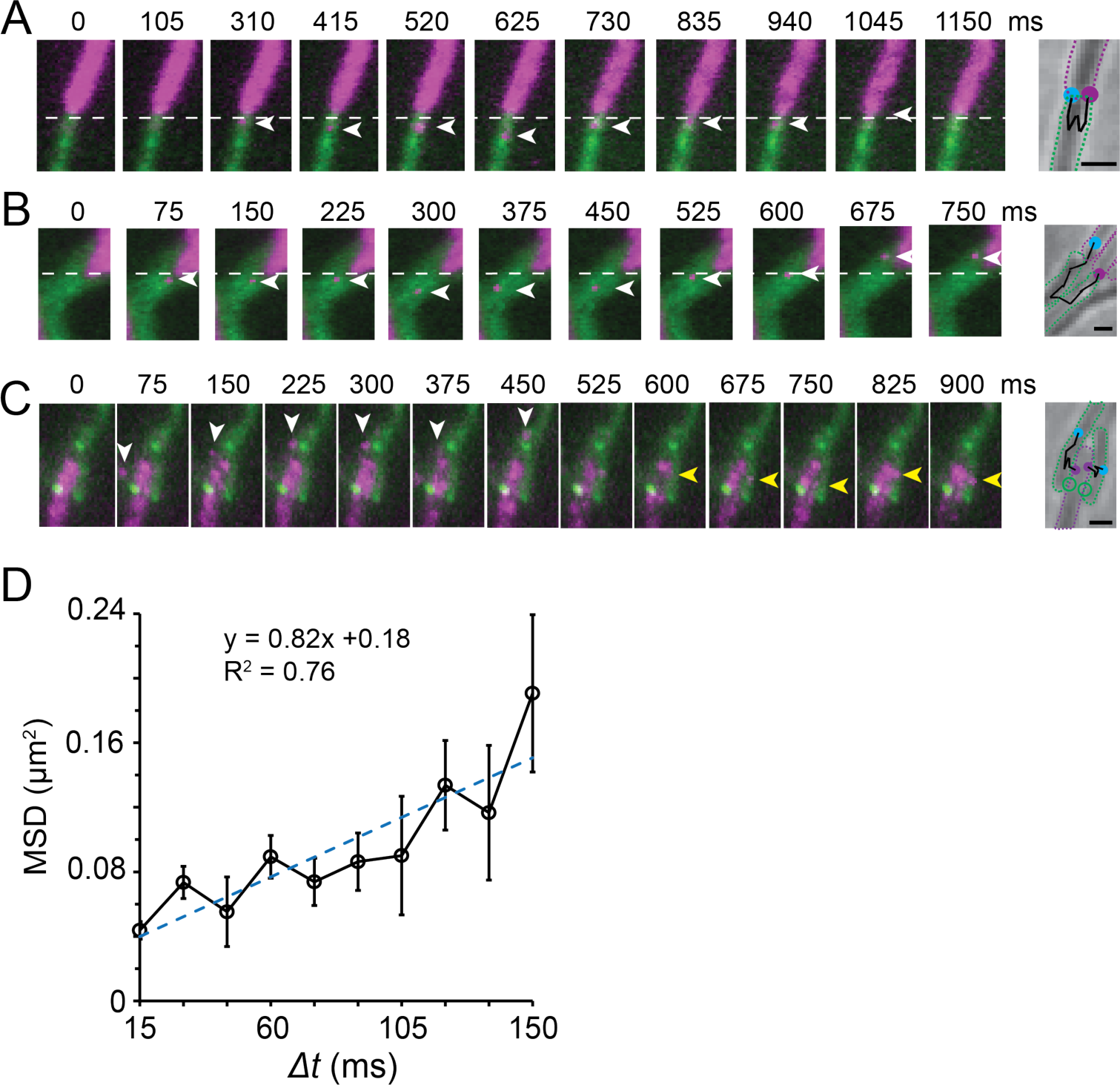
OME requires cargo diffusion. **A - C)** Example trajectories of single SS_OM_-PAmCherry particles during OME. The cells expressing SS_OM_-PAmCherry (magenta) and TraA-GFP (green) were mixed at a 1:1 ratio. Arrows point to the exchanged particles. On the right side of each time-lapse series, the outlines of the donors and recipients are marked with purple and green dotted lines, respectively. The trajectories of particles are marked with black segmented lines. Their start and end positions are marked with purple and blue circles, respectively. Scale bars, 1 μm. The uncropped videos for these OME events are Movies S5, S6, and S7. In A) and B), the initial positions of the donor cell poles are marked with white dashed lines in the time-lapse images. In C), the positions of two TraA foci are marked with green circles on the right panel. **D)** The linear relationship between MSD and *Δt* of the 39 transferred particles indicates they diffuse freely during OME.

By closely tracking particle trajectories during OME, we found six cases where SS_OM_-PAmCherry particles transferred to recipient cells and then returned to the donors (**Fig. 5A, Movie S5**). This observation was consistent with OME occurring bidirectionally, where ‘recipients’ can also act as donors. Strikingly, we identified five SS_OM_-PAmCherry particles being transferred between more than two cells. For instance, the particle shown in **Fig. 5B** and **Movie S6** passed three cell-cell boundaries. Thus, through TraAB-mediated kin recognition, client proteins can be serially transferred between clonemates to achieve long-range OME among multiple adjacent cells.

### TraA foci do not correlate as the channels for OME

Here we sought to investigate the role of TraAB complexes in OME and if they serve as transfer portals. To answer these questions, we followed the trajectories of single-particle OME events between cells that made lateral contacts. Such juxtaposed side-by-side cells frequently formed multiple TraA foci, which allowed us to ask whether these foci correlated as the sites where SS_OM_-PAmCherry particles crossed the cell-cell boundaries. However, out of the seven SS_OM_-PAmCherry particles that transferred through lateral side-by-side cells, none of them passed through the TraA foci (**Fig. 5C, Movie S7**). These observations strongly suggest that TraAB complexes do not form transfer channels for OME.

### A SNAREs-like mechanism for TraAB-mediated OM fusion

To understand how one intercellular TraAB-TraAB dimer induces OM fusion, we used AlphaFold (Jumper *et al*., 2021) to predict the structures of TraA and TraB. Using the sequences of the processed proteins (amino acids 37 – 684 for TraA, excluding its SP and the cleaved MYXO-CTERM (Sah *et al*., 2020) and 23 – 543 for TraB, excluding its SP), AlphaFold generated highly-confident structure models for both proteins in their monomeric states (**Fig. 6A**). TraA monomer forms a dumbbell-like structure where the N-terminal VD, containing at least eight β-sheets, forms the first dumbbell head. The shaft is an extended cysteine-rich region, which makes a 180° turn at the end, forming the second dumbbell head (**Fig. 6A**). This region, containing more than 70 cysteines, forms at least eight β-hairpins that each consists of two antiparallel β-strand connected by a short loop (**Fig. 6A**). We predict that at cell surfaces where thiol groups are oxidized, this region is stiffened by dozens of disulfide bonds. Consistent with a previous finding that the VD determines the recognition between TraA variants (Cao & Wall, 2017), the AlphaFold structure showed two TraA molecules interacting through their VDs, forming a head-to-head dimer (**Fig. 6A**). Strikingly, the length of a TraA dimer is ∼ 28 nm, matching the distance between OMs in tightly packed cell groups (**Fig. 1A**).

**Fig. 6.**
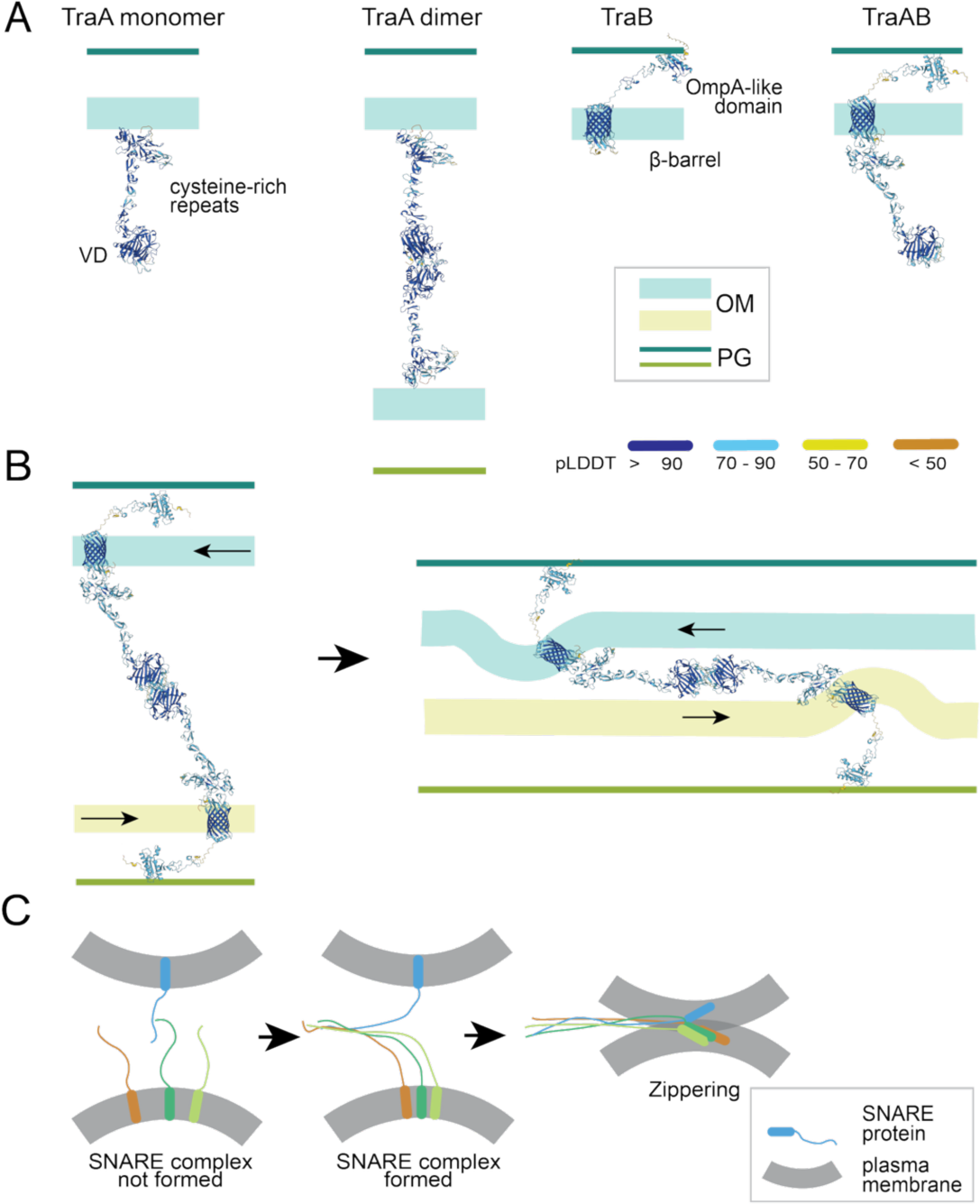
Predicted structures of TraA, TraB, and their complexes suggest a mechanism for OM fusion comparable to the eukaryotic SNARE system. **A)** The predicted structures of TraA, TraB, and their complex. Structure elements are colored according to their pLDDT (predicted local-distance difference test) scores, which indicate the prediction confidence. **B)** A proposed bulge-and-slide mechanism for TraAB-mediated OM fusion. When two cells with compatible TraA molecules make contact, TraA molecules form head-to-head dimers, first through their VDs, and then TraB anchors such TraA dimers to PG. As the juxtaposed cells slide against each other, the two TraAB complexes move in opposite directions, pulling the OMs. Such mechanical stress will bring the OMs of adjacent cells closer and thus trigger OME. **C)** A model of SNARE-mediated plasma membrane fusion in eukaryotic cells. Assembly of SNAREs requires four different SNARE motifs on three or four separate proteins. Once a SNARE complex forms, the four coiled-coil domains transform into a four-helix bundle that “zips” the two membranes together.

The TraB protein consists of a N-terminal β-barrel and a C-terminal OmpA-like domain, linked together by short thrombospondin type 3 (TSP3) repeats predicted to bind calcium (Dai *et al*., 2017) (**Fig. 6A**). Consistent with their colocalization and 1:1 stoichiometry (**Fig. 3**), AlphaFold also predicted a 1:1 TraAB complex in which the second dumbbell head of TraA is positioned on top of the β-barrel in TraB (**Fig. 6A**). However, AlphaFold did not predict significant conformational changes in TraA after its dimerization and interaction with TraB (**Fig. 6A**). Nevertheless, because TraB is diffusive in isolated cells but colocalizes with TraA in the relatively stationary intercellular foci (Cao & Wall, 2019a), TraA dimerization could introduce conformational changes in TraB, which then promotes binding to peptidoglycan (PG) and thus restrains the diffusion of the TraAB-TraAB dimers.

Based on these predictions and our findings, we propose a model for TraAB-mediated OM fusion that has similarities to the eukaryotic soluble N-ethylmaleimide-sensitive factor attachment protein receptors (SNAREs) that mediate vesicle membrane fusion. To facilitate fusion, membranes must be brought close together, and one way to do this is to minimize their contact area. Eukaryotes utilize membrane-anchored SNARE proteins to establish point-to-point contacts that perturb the bilayer structure and thereby lower the energy barrier for fusion. Once SNARE proteins from juxtaposed membranes establish contact, the SNARE complex tighten their coiled-coil domains into bundles, mimicking a zipper that pulls the membranes together until they merge (Jahn *et al*., 2024) (**Fig. 6D**). Through this mechanism, one to three SNARE complexes are sufficient to catalyze membrane fusion (Jahn *et al*., 2024, Sinha *et al*., 2011, Shi *et al*., 2012, van den Bogaart *et al*., 2010). Similarly, we propose that when two cells that express compatible TraA variants approach each other, TraA molecules establish point-to-point contacts between them. Diffusing TraA molecules can then form intercellular dimers in a head-to-head orientation, which is consistent with TraA foci formation between topologically opposed OMs. The TraAB complexes could anchor to PG through the OmpA domain in TraB and thus forms relatively stationary junction foci like SNARE complexes (**Fig. 6B**). While the TraA molecules in a dimer are not likely to form SNARE-like zippers (**Fig. 6D**), when juxtaposed cells slide against each other, the movements of cells pull the PG-anchored TraAB complexes toward opposite directions that tugs on the OM and generates bulges at the base of each complex (**Fig. 6B**). As the cells continue to slide, due to the rigidity of TraA, such stressed OM bulges are primed for membrane fusion as they are forced into juxtaposed OMs, resulting in OM fusion between cells (**Fig. 6B**). In agreement with our model, transmission electron microscopy (TEM) of cryo-fixed thin sections showed that cells overexpressing TraAB contacted each other through OM bulges (Vassallo *et al*., 2015) (**Fig. 6C**).

## Discussion

Multicellular organisms independently evolved from unicellular ancestors at least 25 times during evolution (Grosberg & Strathmann, 2007). Myxobacteria achieve multicellularity through the aggregation of single cells (Kroos *et al*., 2025). While forming aggregates is relatively straightforward, it is less understood how individual cells from genetically mixed populations establish stable multicellular structures. For instance, how do individual cells maintain cooperative cell-cell interactions while keeping cheaters at bay (Velicer *et al*., 2000)? Myxobacteria achieve the transition from solitary, single-cell life into cooperative, multicellular tissues through kin recognition and the exchange of cellular contents (Sah & Wall, 2020). Specifically, the homotypic binding by their polymorphic cell surface TraA protein and its cohort protein TraB, leads to OM content sharing and further discrimination against nonself by OME of polymorphic toxins. To better understand the mechanism of OME we used quantitative microscopy and single molecule tracking. We showed that the ratio of TraA and TraB in cell-cell junctions is 1:1. Moreover, we showed that overexpression of the chromosomal copy of the *traAB* operon generates TraA foci of various stoichiometry, ranging from 2 to 12. Our results confirmed that each partnering cell contributes at least one TraAB complex to form a functional junction. Because the native, low expression of *traAB* is sufficient for OME (Wei *et al*., 2011), we postulate that the minimum number of TraAB complexes in junctions for OME is two. Indeed, the foci that contained two TraAB complexes only accounted for ∼5% of all the foci we detected, which explains why such foci were hard to image when TraAB were expressed with the native promoter (Cao & Wall, 2019a, Vassallo *et al*., 2015). Thus, overexpression increased both the number and size of such foci. The wide range of stoichiometry and the fact that two partnering cells do not necessarily contribute equal numbers of TraAB proteins in foci junctions, suggest that TraAB do not form intercellular protein channels with defined structures.

Instead, our results invoke TraAB serving as fusogens to transiently catalyze the OM fusion between cells and thus allow rapid and facile cargo exchange. This model presumes that the OM of *M. xanthu*s is fluid, distinct from the rigid OMs from other Gram-negative bacteria such as *E. coli* (Cao & Wall, 2020). By tracking single molecules of the lipidated SS_OM_-PAmCherry, we showed this reporter rapidly diffuses in the OM. This analysis thus confirmed *M. xanthus* OM fluidity, showing that diffusion rates in the OM were similar to unrestricted protein diffusion rates in the cytoplasm, IM, and periplasm in other bacteria. It is unclear how widely such fluid OMs exist in Gram-negative bacteria. Nevertheless, our method can be easily applied to measure OM fluidity in other organisms.

Further support of the membrane fusion comes from tracking OME at single-particle resolution. Here SS_OM_-PAmCherry particles were observed to transfer from donor into recipient cells at rates that are nearly identical to their diffusion rates. Strikingly, particles were also observed to enter recipient cells and then return to the original donor, supporting prior observations of bidirectional transfer mediated by OM fusion (Cao & Wall, 2019a). Finally, in tracking experiments, particles also traveled across multiple cell boundaries when those cells were adjacent to one another. This result indicates that multiple cells can simultaneously form fusion junctions, which are in a tissue-like state. This latter point is also supported by prior findings of serial cell-cell transfer of SitA toxins (Vassallo *et al*., 2017).

The last question we addressed is whether TraAB foci serve as transfer portals for OME. However, our observations did not find a correlation between TraA foci and single particle transfer of the SS_OM_-PAmCherry reporter. Instead, transfer occurs at various sites along juxtaposed cells, which are not directly correlated with TraAB foci. Given these findings, we propose TraAB foci serve as sites to nucleate membrane fusion. Once “handshake” recognition occurs between TraAB complexes of juxtaposed cells, a small slide or movement between cells is sufficient to bring the OMs together by pulling forces that generate bulges on OMs, as observed in TEM micrographs of cell cross-sections (Vassallo *et al*., 2015). Additionally, fluorescent time-lapse microscopy of gliding *M. xanthus* cells engaged in OME shows the functional TraA-mCherry reporter dynamically coalescing in a zipper-like fashion along the lengths of the interacting cells (Cao & Wall, 2019a). Taken together, we envision TraAB initiating an OM fusion junction that widens along the cell length. Fission occurs when cells move apart from one another.

The outer leaflet of the *M. xanthus* OM contains LPS, consisting of the lipid A membrane anchor and extended aqueous exposed polysaccharide O-antigen repeats (Perez-Burgos *et al*., 2019). Additional types of lipids may also reside in their outer leaflet, but that is not fully understood (Lorenzen *et al*., 2014). Since TraAB has no known connection to the IM or cytoplasmic energy sources, we propose that when cells slide against each other, the mechanical force of cell movements, an essential requirement for OME (Wei *et al*., 2011), could pull OMs outward through the rigid TraAB-TraAB dimer and eventually cause OM fusion (**Fig. 6B**). Our model shows similarities to the eukaryotic SNARE mechanism in how membranes are brought together to overcome the energy barrier that obstructs membrane fusion (**Fig. 6D**). However, in contrast to the SNARE complexes that require four SNARE motifs from three to four proteins, the TraAB-TraAB dimer is simpler but contains the same essential structural elements: the interaction domains in TraA and the membrane anchor in TraA and TraB (**Fig. 6A**). Different from the SNARE complexes, which pull membranes together through large-scale conformational changes (**Fig. 6D**), the energy for bringing membranes in proximity comes from sliding movements between cells by either motility or cell growth (**Fig. 6B**). Nevertheless, we do not rule out the possibility that conformational changes in TraA could also play important roles in OM fusion. In this regard, a candidate conformational switch within the diverse TraA family is the invariant Ser-Cys-Asn-Cys-Cys-Pro motif found in the C-terminal stalk region (Cao *et al*., 2019). Here the adjacent Cys residues may form a sterically constrained disulfide bond that brakes upon tension by TraA-TraA receptors adhering as cells move in opposite directions.

Our model explains two important observations. First, whereas TraA and TraB do not depend on each other to localize to OMs, their function as cell-cell adhesins and for the formation of junction foci requires both proteins. In this case, TraA is required for kin recognition through its VD to initiate intercellular dimer formation, while TraB is essential to anchor such dimers to PG and to restrain their diffusion. This latter point is supported by the finding that deletion of the OmpA domain from TraB results in a truncated protein able to form foci and act as an adhesion with TraA, but is not functional for OME (Balagam *et al*., 2021). Second, OME requires cell motility (Wall *et al*., 1998, Wall & Kaiser, 1998, Wei *et al*., 2011). Our model predicts that the relative movements pull on OMs through the rigid TraA-TraA dimers, and such mechanical stress will produce pronounced OM bulges that forces OMs together, while also perturbing the bilayer structure, and thereby lowering the energy barrier to the point where fusion occurs spontaneously (**Fig. 6B**). Our model further predicts that after fusion, fission occurs when cells move apart from each other, where long narrow OM tubes are sometimes seen between separating cells (Ducret *et al*., 2013, Wei *et al*., 2014). In future studies, it will be interesting to test if exogenous expression of TraAB in other bacteria with fluid OMs facilitates OME. However, there are technical issues reconstituting the MYXO-CTERM sorting pathway for TraA cell surface localization in a heterologous host (Sah *et al*., 2020, Guo *et al*., 2025). Nevertheless, if successful, this method can be used in synthetic biology for generating bacterial multicellular structures.

## Materials and methods

### Bacterial strains, plasmids and growth conditions

Vegetative *M. xanthus* cells were grown in liquid CYE medium (10 mM MOPS pH 7.6, 1% (w/v) Bacto™ casitone (BD Biosciences), 0.5% yeast extract and 8 mM MgSO_4_) at 32 °C, in 125-ml flasks with vigorous shaking, or on CYE plates that contains 1.5% agar. Strains, plasmids, and primers used in this study are listed in Supplementary **Table 1**. To create P*_pilA_*-*SS_OM_*-*PAmCherry* (pPC60), fragments of P*_pilA_*-*SS_OM_* and *PAmCherry* were PCR amplified (Q5, New England Biolabs), and the resulting amplicons containing 25 bp overhangs were ligated into the pSWU19 vector (linearized with EcoRI and XbaI) in Gibson Assembly Master Mix (New England Biolabs). Construction of the SS_IM_– PAmCherry fusion was done following the published method (Wei *et al*., 2011). This construct was made by PCR amplification of genomic DNA with primers P_pilA_-F2 and SS_IM_–mCherry-R (**Table 1**). The 5′ tail of the latter primer encodes 81 bp from MXAN_1176 that encode the N-terminal IM localization signal plus the downstream 10 amino acids (CKDSDKKESM) for IM retention (Wei *et al*., 2011). This PCR product was cloned to the 5′ end of PAmCherry at the XbaI site. Plasmids were verified by PCR, restriction enzyme digestion, and if necessary, DNA sequencing. Verified plasmids were then electroporated into *M. xanthus* cells and selected with appropriate antibiotics. The strains, plasmids and primers used in this study are listed in **Table 1**.

**Table 1.**
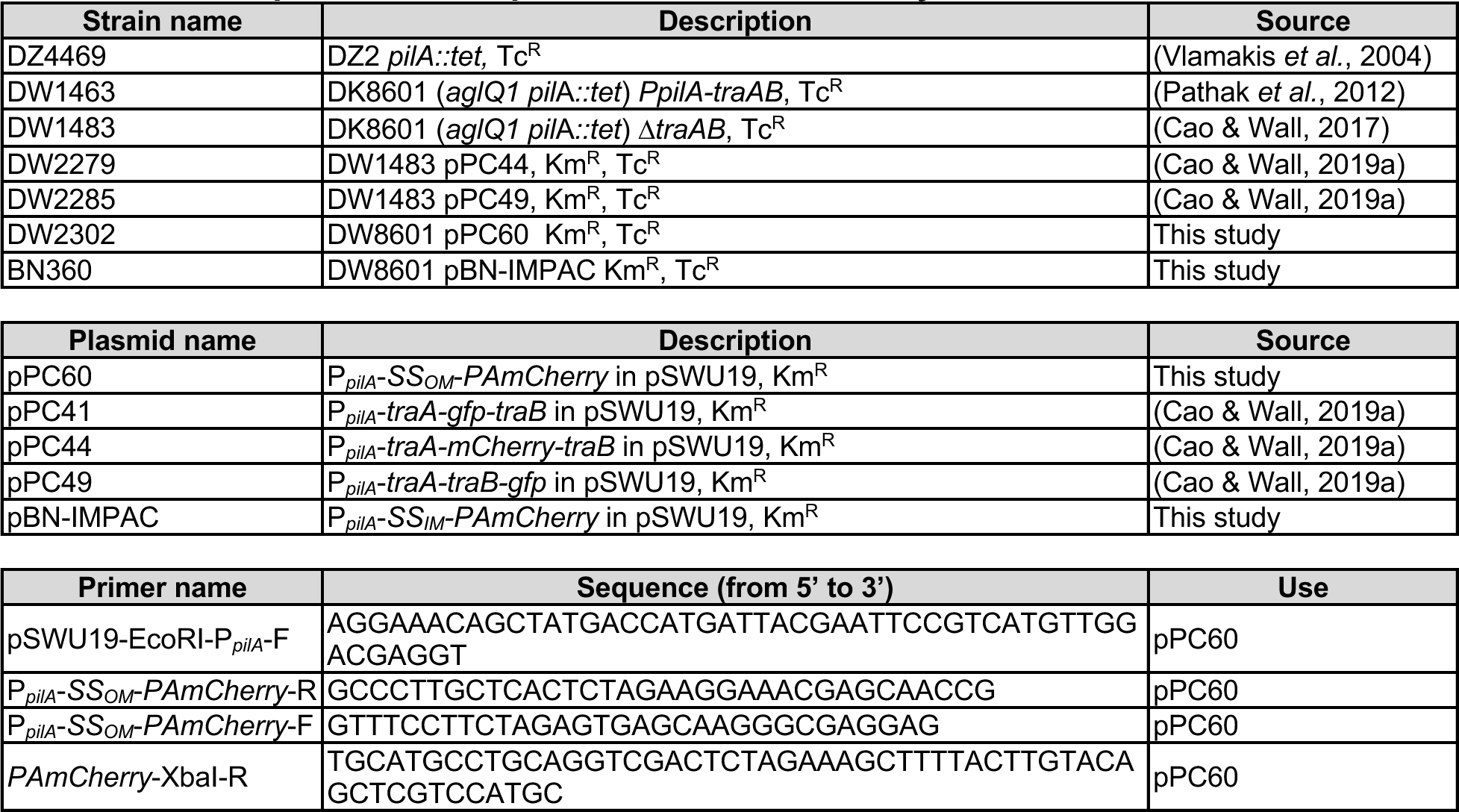
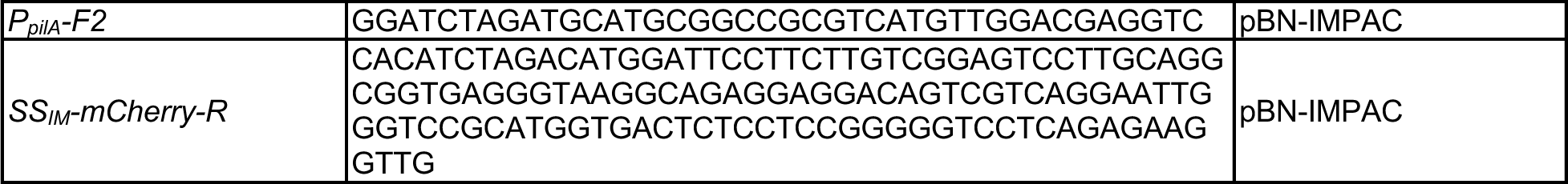
Strains, plasmids, and primers used in this study.

### Light microscopy and image analysis

For all imaging experiments, 5 ml cells or cell mixtures were spotted on a 1.5% (w/v) agar pad containing 10 mM MOPS pH 7.6 and 8 mM MgSO_4_. All microscopy images were captured using an Andor iXon Ultra 897 EMCCD camera (effective pixel size 160 nm) on an inverted Nikon Eclipse-Ti™ microscope with a 100× 1.49 NA TIRF objective. GFP and mCherry were photobleached using the 488-nm and 561-nm lasers (0.5 kW/cm^2^), respectively at the stated frame rates. sptPALM was performed as previously described (Fu *et al*., 2018). *M. xanthus* cells were grown in CYE to OD_600_ around 1. PAmCherry was activated using a 405-nm laser (0.3 kW/cm^2^), excited and imaged using a 561-nm laser (0.2 kW/cm^2^). Images were acquired at 67 Hz (15 ms/frame). For each sptPALM experiment, single PAmCherry particles were localized in at least 100 individual cells from three biological replicates. sptPALM and cell morphology data were analyzed as previously described using a MATLAB (MathWorks) script (Fu *et al*., 2018, Zhang *et al*., 2023a, Zhang *et al*., 2023b). Briefly, cells were identified using differential interference contrast images. Single PAmCherry particles inside of cells were fitted by a symmetric 2D Gaussian function; its center was assumed to be the particle’s position (Fu *et al*., 2018). Particles in consecutive frames were considered to belong to the same trajectory when they were within a user-defined distance of 320 nm (two pixels). Sample trajectories were generated using the TrackMate plugin (Ershov *et al*., 2022) in the ImageJ suite (https://imagej.net). For simplicity, we considered all the mobile particles of these proteins displaying typical diffusion and determined their *D* from a linear fit to the first four points of the MSD using a simpler formula MSD = 4*DΔt* (Fu *et al*., 2018, Lee *et al*., 2016). Error bars were the standard derivation of 1,000 bootstrap samples using the published method (Morgenstein *et al*., 2015).

### Cryo-EM

Bacterial strains were grown overnight in CYE media with appropriate antibiotics, incubated with shaking at 32 °C and 250 rpm to an optical density of OD_600_ 0.6. Cells were collected by centrifugation at 5,000 rpm and room temperature for 3 min and resuspended in CYE media to a final OD_600_ of 12. The cell suspension (3 µl) was applied to C Flat-1.2/1.3 200 mesh copper grids (Electron Microscopy Sciences) that were glow discharged for 30 s at 15 mA. Grids were plunge-frozen in liquid ethane with an FEI Vitrobot Mark IV (Thermo Fisher Scientific) at 4 °C, 100% humidity with a waiting time of 30 s, two-side blotting time of 2.5 - 3 s, and blotting force of 0. All subsequent grid handling and transfers were performed in liquid nitrogen. Grids were clipped onto cryo-FIB autogrids (Thermo Fisher Scientific). We used a Titan Krios G4 transmission electron microscope (Thermo Fisher Scientific), equipped with a Gatan K3 direct electron detector and a Gatan BioContinuum energy filter. Imaging was performed using a 15 eV slit width on the energy filter to enhance image contrast. Micrographs were recorded in counted mode at a nominal magnification of 30,000 x, using a dose rate of 14.65 electrons/pixel/second over a 2.6-second exposure, resulting in a total accumulated dose of 50 electrons/Å². Low magnification images were taken at 6500 x with a pixel size of 1.39 nm.

## Acknowledgments

This work was supported by the National Institutes of Health grants GM129000 to B. N. and GM140886 to D.W.

## Movie captions

**Movie S1.** TraA-GFP foci being photobleached. Time-lapse images were recorded at 67 Hz and played at the same frame rate (real time).

**Movie S2.** Simultaneous photobleaching of TraA-mCherry (right) and TraB-GFP (left) in foci between adjacent cells. Time-lapse images were recorded at 5 Hz and played at the same frame rate (real time).

**Movie S3.** Diffusion of OM_SS_-PAmCherry particles. Time-lapse images were recorded at 67 Hz and played at the same frame rate (real time).

**Movie S4.** Diffusion of IM_SS_-PAmCherry particles. Time-lapse images were recorded at 67 Hz and played at the same frame rate (real time).

**Movie S5**. OME is bidirectional. The donor cell expresses OM_SS_-PAmCherry (magenta) and the recipient TraA-GFP (green). The particle was first transferred to the recipient and then returned to the donor. Time-lapse images were recorded at 67 Hz and played at 7 frames/s, about 1/10 speed of real time.

**Movie S6**. Serial cell-cell transfer of OM_SS_-PAmCherry particle by OME. Donor cell expresses OM_SS_-PAmCherry (magenta) and the recipients TraA-GFP (green). Particle was serially transferred, crossing three cell-cell boundaries. Time-lapse images were recorded at 67 Hz and played at 7 frames/s, about 1/10 speed of real time.

**Movie S6**. TraAB foci do not function as OME conduits. Donor cell expresses OM_SS_-PAmCherry (magenta) and the recipients TraA-GFP (green). Two OM_SS_-PAmCherry particles transferred across cell boundaries, but not through the TraAB foci. Time-lapse images were recorded at 67 Hz and played at 7 frames/s, about 1/10 speed of real time.

